# nf-rare-var-assoc: a comprehensive genetic association pipeline with rich QC

**DOI:** 10.64898/2026.09.19.750380

**Authors:** Piotr Suszyński, Tomasz Gambin

## Abstract

Genetic association analyses of exome sequencing data pose distinct methodological challenges in small cohorts, which are common in rare disease research. Data filtering prior to association testing requires particular care, especially when analyzing datasets generated under different study designs. Most of current genetic association tools need to be integrated into complex data processing pipelines, whose implementation requires technical expertise and is prone to error. To alleviate mentioned issues we built a comprehensive, high performance Nextflow workflow for genetic association testing, which includes all commonly performed data filtering and preparation steps, controlled by a rich set of modifiable parameters. We compared our software against existing tools incorporating quality control and aggregated association testing steps and found our solution to offer the most complete suite of data preparation, association testing and reporting modules, making small-cohort association analyses accessible to teams with limited technical expertise.

## 1. Introduction

Rare-Variant Association Studies (RVAS)(Lee et al., 2014) extend GWAS by increasing statistical power, through variant filtration and aggregation, to detect associations involving rare variants, which may contribute to the missing heritability of complex phenotypes(Rajabli and Kunkle, 2023). Common RVAS methods include burden and SKAT tests(Wu et al., 2011).

Genome sequencing data is particularly valuable for rare variant association testing. Modern variant callers, such as DeepVariant(Poplin et al., 2018) deliver individual genotype likelihoods, which can be used to calculate soft-genotypes, dosages, further enriching the data. While there is no lack of software tools for harnessing the sequencing data for RVAS, these tools need to be integrated into complex data processing pipelines, executing data normalization, multiple steps of filtering, with branching data flows for kinship analysis and filtering, LD-based filtering, PCA, fitting population structure models. Unfortunately, most of the pipelines currently available are either still tailored to traditional genotyping-array-based GWAS, having different quality control (QC) requirements, or offer only rudimentary QC procedures. As it turns out, there *is* a lack of comprehensive end-to-end pipelines, including both QC and association testing, tailored to genome sequencing data.

To address these limitations and tap into the wealth of data offered by Genome or Exome Sequencing (ES), particularly in the context of rare-disease small-cohort studies, we developed the *nf-rare-var-assoc* Nextflow(Di Tommaso et al., 2017) pipeline, which includes all the data filtering and preparation steps commonly performed before running the tests, as well as reporting and some less common steps that we found to be beneficial, such as dosage calculation and using it as soft-genotypes for association testing. The pipeline is highly configurable, enabling the user to adapt the filtering thresholds and parameters controlling various processing steps, as well as toggle selected subsets of functionalities.

In our work we are not limiting ourselves to rare variants only. Instead we focus on “small cohorts” characteristic-with samples count below 3000. For certain small cohorts it might be possible even for common variants to have relatively large effect sizes. An example may be the task of searching for genetic grounds responsible for subphenotypes within rare disease cohorts such as congenital heart defects occurring in 60-80% of individuals with 22q11.2 deletion syndrome(Goldmuntz, 2020; Smyk et al., 2023), which can be viewed as a very common phenotype within this cohort. Thus even common variants, when co-occurring with the main deletion, can potentially have large effects. At the same time, the very small sample sizes available for such conditions justify the use of RVAS techniques, in order to increase the statistical power of the tests.

### Summary of contributions

- an automated pipeline for rare-variant association testing which includes a rich set of data preparation/QC steps
- dosage calculation from PL values which are estimated from GQ if missing, with multiallelic sites correction, and incorporation of dosage handling into the whole pipeline
- a DataFusion query optimization which enables processing genotype data in polars-bio(Wiewiórka et al., 2025) using SQL with lower resource requirements
- review and comparison of quality control and association testing pipelines

## 2. Methods

The pipeline ingests a multi-sample vcf file with genotype (GT), genotype quality (GQ), coverage depth (DP) and optionally dosage (DS) or Phred-scaled genotype likelihoods (PL) fields. The second input is the phenotype, which can be either two files listing cases and controls or a single tab-delimited file with a phenotype column. The pipeline consists of the following configurable steps (see Figure 1a):

**Figure 1:**
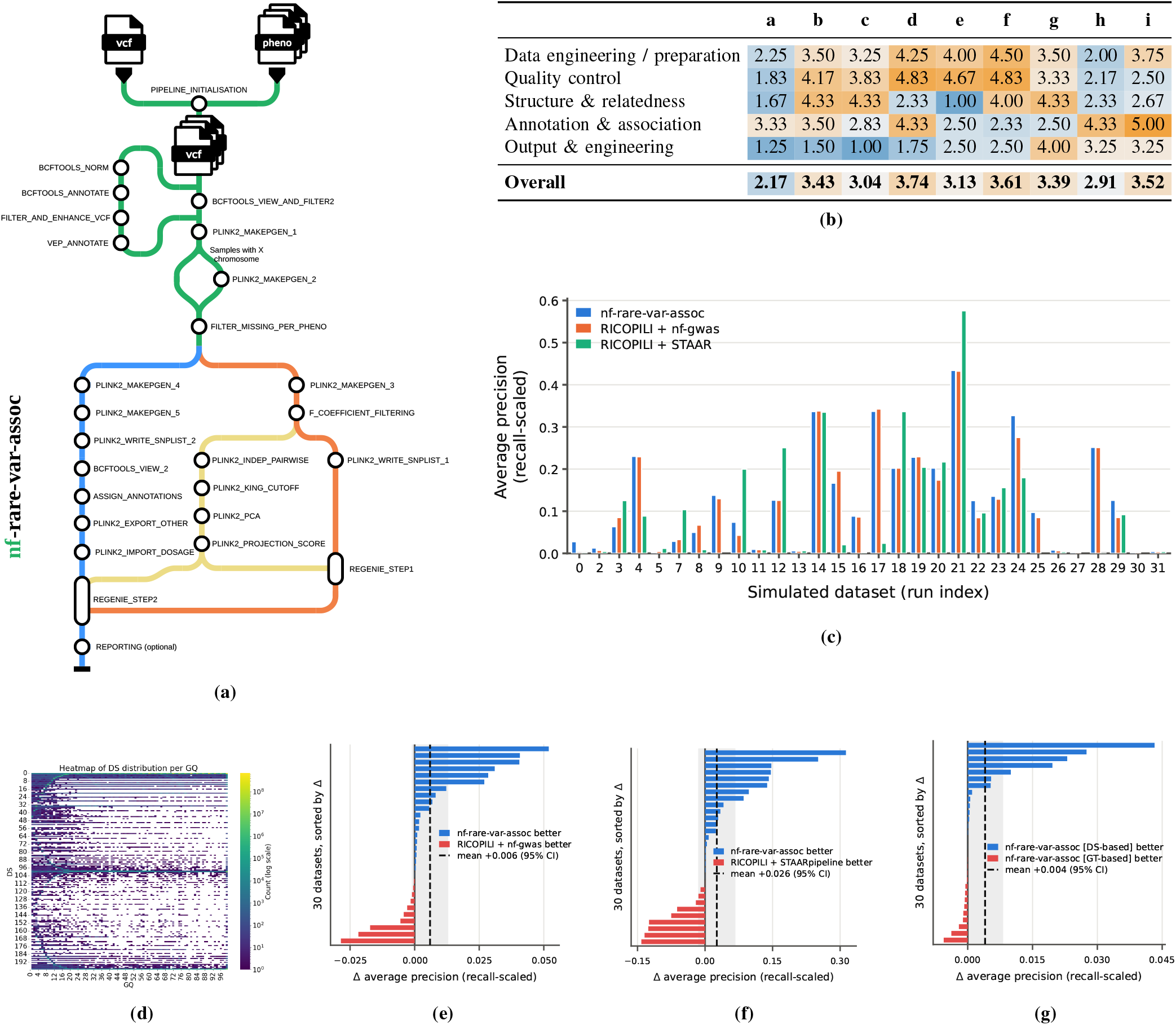
**(A)** Diagram of the nf-rare-var-assoc pipeline. **(B)** Functionality comparison of nf-rare-var-assoc to similar association pipelines and QC-focused pipelines. Averages of dimensions graded 1–5 by completeness, lower = more complete. Full comparison, along with details on each dimension and the grading criteria, available in appendix D. Compared tools: **a**: nf-rare-var-assoc, **b**: genepi/nf-gwas(Schönherr et al., 2024), **c**: HTGenomeAnalysisUnit/nf-pipeline-regenie(Giacopuzzi, 2026), **d**: montilab/nf-gwas-pipeline(Song et al., 2021), **e**: UW-GAC GENESIS analysis pipeline(Gogarten et al., 2022), **f**: STAARpipeline(Li et al., 2022), **g**: SEQSpark(Zhang et al., 2017), **h**: RICOPILI(Lam et al., 2020), **i**: plinkQC(Syed et al., 2025). **(C)** Empirical gene-detection performance on the simulated benchmark datasets: recall-scaled average precision per dataset for nf-rare-var-assoc, RICOPILI + nf-gwas and RICOPILI + STAAR. RICOPILI + STAAR: didn’t work for chromosome X, failed to produce results for datasets 9, 25, 27, 30. The average precision for datasets 0, 16, 28 for RICOPILI + STAAR was below 3 10^*−*3^ and is not clearly visible on the plot, even though it is non-zero. The same is true for datasets 5, 27 and 30 for nf-rare-var-assoc and for datasets 0, 27, 30 for RICOPILI + nf-gwas. **(D)** One of the plots generated during dataset characteristics analysis in nf-rare-var-assoc, showing the distribution of dosage (DS) as a function of genotype quality (GQ). **(E)** The RICOPILI + nf-gwas pairwise comparison: per dataset differences in recall-scaled average precision. nf-rare-var-assoc leads on 20/30 datasets; the mean paired difference is +0.00596, 95% CI [*−*0.00076, 0.01268] (dashed line, shaded band) - not significant (paired *t* test, *p* = 0.0798). **(F)** The RICOPILI + STAARpipeline pairwise comparison: per dataset differences in recall-scaled average precision. nf-rare-var-assoc leads on 20/30 datasets; the mean paired difference is +0.02613, 95% CI [*−*0.01335, 0.06560] (dashed line, shaded band) - not significant (paired *t* test, *p* = 0.1863). **(G)** nf-rare-var-assoc dosage ablation pairwise comparison: per dataset differences in recall-scaled average precision. nf-rare-var-assoc with dosage use leads on 16/30 datasets with 1 tie; the mean paired difference is +0.00402, 95% CI [0.00006, 0.00798] (dashed line, shaded band) - significant (paired *t* test, *p* = 0.0470).

1. We split multi-allelic sites, left-align and normalize, remove duplicated variants, and assign variant ids.
2. We select a subset of samples, filter variants by `QUAL`, average `GQ` and average `DP` values, set genotypes of individual samples to missing value.
3. We use VEP(McLaren et al., 2016) to annotate variants.
4. To preserve as much data as possible we employ a multi-step missingness filtering approach, with separate variants and samples-based filtering along with per-phenotype variants missingness filtering, which is targeting data with differences in cases and controls missingness, for example caused by different sources and study designs of ES data. Optionally different filtering thresholds for data flowing to REGENIE(Mbatchou et al., 2021) step 1 and step 2 can be used.
5. We infer sex from the X chromosome data. It can be included in covariates passed to REGENIE.
6. If absent, we calculate the dosage using the `PL` field, which in real data may be missing for genotypes 0/0 - therefore we first fill in the lacking `PL` values by estimating them from genotype quality `GQ`. We also apply multiallelic-sites-specific correction. To achieve high performance and low memory use we used polars-bio with custom query optimization implemented by us. For details please see appendix A and B.
7. Before PCA and REGENIE step 1, we filter variants based on minor allele count, minor allele frequency, and deviations from Hardy-Weinberg equilibrium.
8. At the same time we also filter samples with too high or too low inbreeding coefficient - we remove outliers using the number of standard deviations parameter.
9. By default, kinship filtering is applied only to the data used for PCA.
10. We have separate PLINK(Chang et al., 2015) LD filtering for the `PCA` path and the `F_COEFFICIENT_FILTERING`, which allows to use different thresholds for those modules. Apart from that for `PCA` we also remove known high-LD genome regions, configured with a file.
11. We compute the PCA and then include top N principal components as covariates in REGENIE step 1 and 2.
12. REGENIE supports grouping variant annotations into masks and executing tests for those masks. We select annotations produced by VEP by severity to assign a mask key and save the data in formats expected by REGENIE.
13. Genomic association testing is done by REGENIE, which supports aggregation methods such as Burden tests, SKAT and SKAT-O(Lee et al., 2012).
14. We produce a single HTML report containing Manhattan plots, Q-Q plots as well as a number of plots describing the characteristics of the data, including PCA projection plot, distributions of GQ, DP and dosage for variants and for samples as well as per phenotype (see example in Figure 1d).

## 3. Comparison to existing tools

### 3.1. Functionalities comparison

We compared *nf-rare-var-assoc* to other tools offering similar capabilities or their subsets. We ranked each tool on 23 features/capabilities - the full ranking is available in appendix D and Figure 1b contains an aggregated version, with capabilities grouped to 5 categories. The details of the grading criteria and justifications for each score can be found in appendix D.

The comparison highlighted unique capabilities of the *nf-rare-var-assoc* pipeline, which is the only end-to-end automated pipeline with rich QC procedures, robust structure and relatedness modeling and featuring association testing tailored to small cohorts and rare diseases. The pipeline also includes comprehensive input preparation steps, allowing it to work on VCF files, without the need to manually process and normalize them.

### 3.2. Evaluation on simulated datasets

To validate the software we implemented a high quality set of automated nf-test based unit tests and executed an empirical evaluation. On a subset of 1000 Genomes Project(Byrska-Bishop et al., 2022) data (exome of chromosomes 12, 22 and X) we simulated 30 diverse phenotypes and ran association testing with *nf-rare-var-assoc*, followed by evaluation where we compared the results with the ground truth used during phenotypes generation. This allowed us to calculate the area under the precision-recall curve for each dataset. Since no competing tool spans the range of quality control and association testing capabilities which *nf-rare-var-assoc* provides, we assembled two chains for comparison, consisting a combination of QC and PCA calculation done with *RICOPILI* followed by association testing done with either *nf-gwas* or the *STAARpipeline*. The latter, contrary to the name, is not an orchestrated pipeline but an R library, lacking advanced QC procedures. Using this tool required us to develop complex R and shell scripts to invoke external tools such as PLINK(Chang et al., 2015) and BCFtools(Danecek et al., 2021) and perform data conversions using sed and awk. Together, these scripts comprised more than 1,400 lines of code (see appendix C). This highlights one of the biggest benefits of using *nf-rarevar-assoc* - the same outcome was achieved with just a single Nextflow call.

The recall-scaled average precision for these comparisons can be seen on Figure 1c. Pairwise differences shown on Figures 1e and 1f indicate that *nf-rare-var-assoc* was better at recovering true positives while inhibiting false positives than the next best solution, which was *RICOPILI + STAARpipeline*, but the difference was not statistically significant with the number of datasets we used. The main advantage of *nf-rare-var-assoc* we want to highlight remains the end-to-end automation of both QC and small sample size targeted association testing.

## 4. Conclusion

We presented *nf-rare-var-assoc*, an all-in-one Nextflow pipeline targeted for small samples size association testing, incorporating rich set of quality control and data preparation steps, enabling easy association testing execution. In our feature comparison the *nf-rare-var-assoc* came out as the only available tool incorporating both rich QC and aggregation-based association testing and at the same time being a highly automated pipeline rather than a software library or a toolkit. Our empirical evaluation indicates the more advanced data filtering and dosage use might lead to increased efficacy of subsequent association testing stages.

## 5. Statements

### Conflicts of interest

The authors declare that they have no competing interests.

## Funding

Work on this project was financially supported by the Warsaw University of Technology grant WUM PW INTEGRA-2/8/2026.

We gratefully acknowledge Polish high-performance computing infrastructure PLGrid (HPC Center: ACK Cyfronet AGH) for providing computer facilities and support within computational grant no. PLG/2025/018863.

## Data availability

The Thousand Genomes data is publicly available and the generated phenotypes are available, along with all the source code, in the https://github.com/biodatageeks/nf-rarevar-assoc git repository.

## Author contributions statement

P.S. implemented the nf-rare-var-assoc pipeline, compared functionalities to competing tools, conducted experiments on Thousand Genomes data and wrote the manuscript.

T.G. implemented an initial proof of concept code, supervised the work on nf-rare-var-assoc and the experiments, reviewed and edited the manuscript.

## Artificial Intelligence tools use

AI tools were used for the source code and project documentation writing, for researching similar methods and for reviewing the source code of the compared tools for the purpose of functionalities comparison.

## Appendix A.

### Dosage calculation

If it is missing, we calculate dosage (`DS` field) from the Phred-scaled genotype likelihoods (`PL` field) by converting to genotype likelihoods 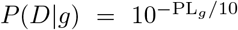 and then observing that the genotype probability can be obtained using the Bayes formula:

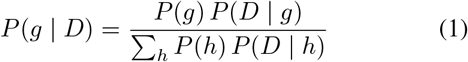

and the dosage is given by:

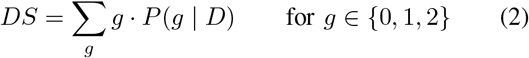

We derive the prior from the allele frequencies *p* and by default we apply a Hardy-Weinberg prior, which for haploid sites takes the form of *P* (*g* = 0) = 1*− p, P* (*g* = 1) = *p* and for diploid sites we have:

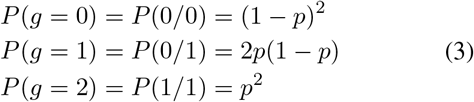

#### A.1. Handling malformed PL values

However, we observed that PL values in VCF files produced by the DeepVariant variant caller already contain the posteriors, so we try to detect the variant caller that was used to produce the VCF file and for DeepVariant we employ a flat prior: 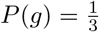 for diploid and 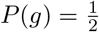for haploid.

Second observation for DeepVariant + GLNexus produced VCF files is that for genotypes 0/0 we observed that a fraction of data may carry all-zeros `PL` field, even when `GQ` is non-zero, which prevented us from calculating dosage. To overcome this problem we estimated the probability of the called genotype being incorrect for such cases from the `GQ` field by assigning:

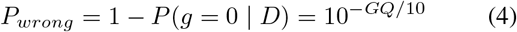

This matches the VCF specification of the `GQ` field meaning, which DeepVariant respects. GATK variant caller uses a different definition of `GQ` but it doesn’t matter for us since we never observed all-zeros `PL` values produced by GATK.

We then need to split this probability between *P* (*g* = 1| *D*) and *P* (*g* = 2 |*D*) and we do that in proportion to their prior probabilities:

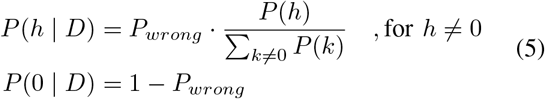

Then, for a diploid ‘0/0’ call with the Hardy-Weinberg prior this gives:

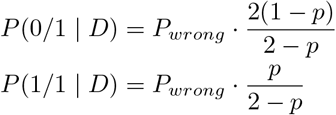

and the final dosage:

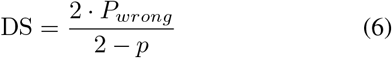

for haploid sites DS = *P*_*wrong*_.

It must be noted that this step runs only for genotypes 0/0 (or 0 in haploid case) with missing `PL`. The biggest impact is for for low `GQ` sites, when *P*_*wrong*_ is non-negligible - distributing this probability mass results in more information about the genotype reaching association tests instead of being ignored, as is the case for the hard genotypes.

#### A.2. Handling multiallelic sites

Multiallelic sites present additional challenges. We used bcftools to split such sites to biallelic, but this resulted in genotypes whose PL field value accounts for just a fraction of the genotype probability mass for some samples. Let’s use a simple example to illustrate the issue. Suppose that there is a VCF record with four alleles, REF=A and ALT=C,G,T and a sample whose genotype is C/G. After the split we could end up with VCF rows presented in Table S1 (showing GT and PL values only for the mentioned sample, in reality this is a multi-sample VCF).

**TABLE S1:** Result of splitting a multiallelic site.

| REF | ALT | GT | PL |
| --- | --- | --- | --- |
| A | C | 1/0 | 300,40,12 |
| A | G | 0/1 | 300,44,14 |
| A | T | 0/0 | 300,260,247 |

In all those rows the minimum PL value is not 0 and in the case of the last row it is very large, because the sample actually doesn’t have both the ALT allele *nor* the REF allele. For the first two rows the PL values encode the fact that the sample has some of the ALT allele but none of the REF allele. A naive application of Equations 1 and 2 gives a dosage value very close to 2 for all those records, which incorrectly describe the genetic reality of the sample.

To overcome such issues, we decided to set DS to missing when the lowest PL value is higher than a given threshold `ds_max_pl_min` (we used 30 as the default value after checking how different thresholds affect the difference between the aggregated DS and aggregated GT values).

Additionally, instead of Equation 2 use the following Equation for final dosage calculation:

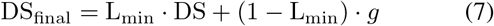

where DS is the dosage calculated using Equation 2, 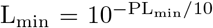 and *g* is the ALT allele count, based on the GT value. When the lowest PL value is 0, as is the case for the vast majority of records, L_min_ is 1 and Equation 7 recovers the original dosage calculation. When *<* PL_min_ *<* `ds_max_pl_min` the (1 *−*L_min_) *· g* term pulls the dosage to the hard-genotype, weighted by a term describing the lack of information in the PL field.

#### A.3. Workaround due to problems with plink

Exporting to traw^1^ file and importing just before association testing was necessary because not all tools and operations currently support dosage information. Since we encountered problems with sex imputation and heterozygosity calculation when dosage was imported into pgen/pvar/psam files, we decided to execute these steps without dosage data and import it just before the REGENIE steps.

#### A.4. Ablation for dosage use in association testing

During our evaluation on simulated datasets (see appendix C) we conducted experiments for nf-rare-var-assoc with and without dosage use, calculated as described above. We present those results in Table S2. Dosage use was statistically significantly improving performance and is responsible for a part of the lead that nf-rare-var-assoc enjoyed in our experiments over compared solutions.

## Appendix B.

### unnest *→*transform *→*array agg DataFusion query optimization

Calculating the dosage field (DS) from the genotype likelihoods field (PL) with filling missing values by genotype quality (GQ) derived estimate proved to be computationally demanding using available tools. We first implemented the transformation using the pysam, and then cyvcf2 libraries, but were not satisfied with the running time. We reimplemented the transformation using the Rust-Htslib library, which improved the running time but resulted in the logic of the transformation being hardcoded into an executable built into a docker container, which means it was invisible from the Nextflow pipeline user perspective. In the end, we decided to switch to an implementation basing on the polars-bio library, which allowed us to express the whole transformation in SQL, visible to the Nextflow user and easily modifiable. It also allowed us to process the VCF in a streaming fashion, limiting peak memory usage, as never all the data needs to be loaded into the memory.

polars-bio is based on the DataFusion execution engine and contains genetics-specific extensions, including VCF support. The genotype fields, such as GQ, DS or PL, are stored, for each variant, in a `genotypes` structure which contains an array per genotype field. In each of those arrays we have the values for consecutive samples, so, in the case of the data we used, each array was of length 3202. Calculation of PL basing on GQ and GT, and also calculation of DS basing on PL, requires to access the arrays elements for corresponding samples. In SQL this can be done by a query of the form:

**TABLE S2:** Ablation for the nf-rare-var-assoc: using dosage versus not using it. A pairwise comparison difference test for average precision (AP) gave *p*_*value*_ = 0.047 for the “use dosage” being better.

| Dataset | no-dosage AP | dosage AP | AP difference |
| --- | --- | --- | --- |
| 0 | 0.033333 | 0.027778 | 0.005556 |
| 2 | 0.013410 | 0.012804 | 0.000606 |
| 3 | 0.063260 | 0.063355 | -0.000094 |
| 4 | 0.202815 | 0.230307 | -0.027492 |
| 5 | 0.001905 | 0.001909 | -0.000004 |
| 7 | 0.023348 | 0.028656 | -0.005308 |
| 8 | 0.051007 | 0.049903 | 0.001104 |
| 9 | 0.137999 | 0.137926 | 0.000073 |
| 10 | 0.074946 | 0.073935 | 0.001011 |
| 11 | 0.010439 | 0.009647 | 0.000792 |
| 12 | 0.126368 | 0.126314 | 0.000053 |
| 13 | 0.005653 | 0.006076 | -0.000423 |
| 14 | 0.336340 | 0.336462 | -0.000122 |
| 15 | 0.123473 | 0.166710 | -0.043237 |
| 16 | 0.087552 | 0.088559 | -0.001006 |
| 17 | 0.337197 | 0.337009 | 0.000188 |
| 18 | 0.201581 | 0.202076 | -0.000496 |
| 19 | 0.230271 | 0.228231 | 0.002040 |
| 20 | 0.192387 | 0.202366 | -0.009979 |
| 21 | 0.434664 | 0.434128 | 0.000536 |
| 22 | 0.105712 | 0.125312 | -0.019599 |
| 23 | 0.130161 | 0.135571 | -0.005410 |
| 24 | 0.326923 | 0.326923 | 0.000000 |
| 25 | 0.101042 | 0.097258 | 0.003784 |
| 26 | 0.008708 | 0.008033 | 0.000675 |
| 27 | 0.001656 | 0.001548 | 0.000108 |
| 28 | 0.251168 | 0.251506 | -0.000338 |
| 29 | 0.102904 | 0.125925 | -0.023021 |
| 30 | 0.001560 | 0.001568 | -0.000008 |
| 31 | 0.004096 | 0.004683 | -0.000587 |

~~~
**WITH indexed AS** (
**SELECT** *,
ROW_NUMBER() OVER () **AS** variant_idx
**FROM** vcf_table
),
samples_unnested **AS** (
**SELECT** variant_idx, UNNEST(genotypes.”GQ”) **AS** gq, UNNEST(genotypes.”PL”) **AS** pl
**FROM** indexed
),
transformed **AS** ([…])
**SELECT**
variant_idx, STRUCT(
array_agg(gq) **AS** GQ, array_agg(corrected_pl) **AS** PL, array_agg(calculated_ds) **AS** DS
) **AS** genotypes **FROM** transformed **GROUP BY** variant_idx
~~~

The actual calculation is performed in the `transformed` section, where we have access to GQ and PL values of the corresponding samples.

However, it turned out that executing such a query caused an explosion in memory usage, consuming tens of gigabytes for even a relatively small, filtered VCF and causing the process to crash by using up all the system memory. Clearly, DataFusion engine couldn’t process the data in a streaming way and tried to load the whole file into the memory.

To fix the memory issue, we implemented a custom DataFusion logical optimizer rule and physical execution plan implementation, called FusedArrayTransform. The logical rule detects queries of the form shown above (unnest *→*transform *→* array_agg) and for those queries executes our custom optimized memory-bounded streaming transformation. It supports arbitrary transformations of the unnested data and is well suited for efficient VCF genotype fields handling.

Our optimizations were submitted as Pull Requests:

- datafusion-bio-functions repository: https://github.com/biodatageeks/datafusion-bio-functions/pull/31
- polars-bio repository: https://github.com/biodatageeks/polars-bio/pull/349

## Appendix C.

### Evaluation on simulated datasets

#### C.1. Selection of the 1000 Genomes subset

We downloaded vcf.gz files for chromosomes 12, 22 and X from http://ftp.1000genomes.ebi.ac.uk/vol1/ftp/data_collections/1000G_2504_high_coverage/working/20201028_3202_raw_GT_with_annot^2^. These are 1000 Genomes Project data, processing of which is described in detail in http://ftp.1000genomes.ebi.ac.uk/vol1/ftp/data_collections/1000G_2504_high_coverage/working/20201028_3202_raw_GT_with_annot/20201028_1000G_2020Oct26_NYGC_JointGenotyping_README.pdf

In order to reduce the file sizes we filtered them to only exome regions and also removed all variants with average genotype quality lower than 4. Below is the script that was used (showing for only one of the chromosomes):

~~~
podman run --rm -v $PWD/:/wd/:z quay.io/ biocontainers/bcftools:1.20--h8b25389_0 bcftools filter --output /wd/20201028
_CCDG_14151_B01_GRM_WGS_2020-08-05_chr12.
recalibrated_variants.exome.vcf.gz --regions-file /wd/TwistRefseqClinvarTarget_hg38_basic. bed --exclude ‘AVG(FORMAT/GQ)<4’ --output-**type**
z --write-index=tbi --threads 4 /wd/20201028
_CCDG_14151_B01_GRM_WGS_2020-08-05_chr12.
recalibrated_variants.vcf.gz
podman run --rm -v $PWD/:/wd/:z quay.io/ biocontainers/bcftools:1.20--h8b25389_0 bcftools concat --allow-overlaps --output /wd
/20201028_CCDG_14151_B01_GRM_WGS_2020-08-05
_chr_12_22_X.recalibrated_variants.exome.vcf. gz --output-**type** z --write-index=tbi --threads
4 /wd/20201028_CCDG_14151_B01_GRM_WGS_2020
-08-05_chr12.recalibrated_variants.exome.vcf. gz /wd/20201028_CCDG_14151_B01_GRM_WGS_2020
-08-05_chr22.recalibrated_variants.exome.vcf. gz /wd/20201028_CCDG_14151_B01_GRM_WGS_2020
-08-05_chrX.recalibrated_variants.exome.vcf.gz
podman run --rm -v $PWD/:/wd/:z quay.io/ biocontainers/bcftools:1.20--h8b25389_0 bcftools sort --output /wd/20201028
_CCDG_14151_B01_GRM_WGS_2020-08-05_chr_12_22_X
.recalibrated_variants.exome.sorted.vcf.gz --output-**type** z --write-index=tbi /wd/20201028
_CCDG_14151_B01_GRM_WGS_2020-08-05_chr_12_22_X
.recalibrated_variants.exome.vcf.gz
~~~

#### C.2. Construction of the simulated datasets

We simulated 30 phenotypes on the real genotype data from 1000 genomes project. We executed following steps:

1. **Variant filtering**. Variants were annotated with Ensembl VEP and then filtered on:
  - the VEP consequence field `CSQ`
  - site `QUAL` and per-site average `FORMAT/GQ` and `FORMAT/DP`
  - a minor-allele-frequency window
  - a maximum allele length, which bounds the size of indels considered
  The filtering thresholds were randomly selected per dataset as given in Table S3.
2. **Selection of causal genes and variants**. From the genes that still carried at least `min_variant_count` variants after filtering, `k_genes` genes were drawn at random; within each gene, a random fraction `variant_fraction` of its remaining variants was designated as causal. The selected genes and the causal variants are the truth against which all the tested tools/pipelines are scored.
3. **Phenotype simulation**. A binary phenotype was simulated using GCTA(Yang et al., 2011) from those causal variants using:

~~~
gcta64 --simu-cc <n_case> <n_control> --simu-hsq <h2> --simu-k <K> --simu-rep 1 --simu-causal-loci <snplist_file> --bfile <input>
~~~

The requested counts were always `--simu-cc 667 1333` (a cohort of 2,000 at a 2:1 control-to-case ratio). The number of controls remained fixed at 1,333 in every dataset, while the number of cases varied between 11 and 667 (median 117), depending on the disease prevalence parameter *k*.

Table S3 lists the sampling domain of every parameter. Phenotype files and causal-truth files for all thirty datasets are distributed with the pipeline repository^3^, so the benchmark can be repeated without re-running the simulation. The datasets span a wide range of difficulty: 1 to 12 causal genes (median 6) carrying 2 to 196 causal variants (median 41) and trait heritability between 0.036 and 0.699 (median 0.413). 6 out of 30 datasets were generated without filtering prospective causal variants with annotations so low impact variants, such as synonymous variants, might have been selected as causal in those datasets. This makes the datasets challenging, and it is expected that for some of them it might be impossible to find the causal variants or genes with the setup of those datasets.

#### C.3. Running nf-rare-var-assoc

nf-rare-var-assoc supports running it for multiple phenotype files (for a single multi-sample VCF file) in parallel and this was utilized in our experiments. VCF preparation steps, including normalization, variant-identifier assignment, VEP annotation and dosage computation from genotype likelihoods - depends only on the genotypes and not on the phenotype, so they were run once and the preparation output was then reused with `--skip_preparation true` flag. We used following parameters:

~~~
use_dosage: true filter_vcf_qual_min: 23
filter_vcf_avg_gq_min: 23
filter_vcf_avg_dp_min: 23
filter_vcf_avg_dp_max: 110
filter_vcf_sample_gq_min: 11
filter_vcf_sample_dp_min: 22
filter_vcf_sample_dp_max: 340
inbreeding_outliers_range_stds: 6 plink2_makepgen_3_options: “--geno 0.25 --hwe 1e-9
0.01 --mac 15 --maf 0.05”
plink2_makepgen_4_options: “--geno 0.1”
plink2_makepgen_5_options: “--mind 0.2”
plink2_write_snplist_qc_options: “--mind 0.15”
plink2_indep_pairwise_options: “--mind 0.3”
plink2_indep_pairwise_window: “800 80 0.2”
plink2_missing_per_pheno_options: “--geno 0.35”
plink2_indep_pairwise_window_pca: “800 80 0.2”
plink2_king_cutoff_threshold_pca: 0.19
plink2_write_snplist_step2_options: “--mind 0.2” regenie_step1_options: “--bt --bsize 400 --
covarColList PC1_AVG,PC2_AVG,PC3_AVG,PC4_AVG” regenie_step2_options: “--bt --minMAC 7 --ref-
first --firth --approx --bsize 200 --aaf-bins
0.1 --vc-tests skato --covarColList PC1_AVG, PC2_AVG,PC3_AVG,PC4_AVG”
~~~

These values were selected basing on our prior experiments on 1000 genomes data, conducted before the testing datasets used in the currently described experiments were generated. To understand how each of those parameters is used in nf-rare-var-assoc please consult Table S4 for the mapping between parameter names and the module names shown on Figure 1A from the main paper.

The association results produced by the pipeline were then scored against ground truth causal genes by calculating average precision for each dataset. The produced metric was then multiplied by the number of causal genes present in the association results divided by the total number of causal genes used for generating the dataset (recall scaling) - this was done to penalize pipelines for filtering too much data which might filter variants for certain genes completely.

The exact commands that we used are available at:

~~~
docs/tool-comparison/benchmark-common/ run_nf_rare_var_assoc.sh
~~~

in the git repository. The whole processing is handled by a single nextflow call and another one for the results scoring.

#### C.4. Running RICOPILI + nf-gwas

We used RICOPILI for quality control and PCA calculation and projection followed by nf-gwas (with the QC available in it) executing REGENIE. Neither RICOPILI nor nf-gwas can produce the masks files for variant grouping association testing so we re-used the ones (annotations and setlist files) produced by the nf-rare-var-assoc pipeline - this was the only thing “loaned” from nf-rare-var-assoc. We tried to use QC parameters as similar as the ones used for the nf-rare-var-assoc run as possible, whenever a corresponding step was present in either RICOPILI or nf-gwas.

The complete script running the RICOPILI + nf-gwas pipeline is available in:

~~~
docs/tool-comparison/chained-benchmark-nf-gwas/ run_chain_ricopili_nf_gwas.sh
and the RICOPILI docker image was created using Dock-erfile available at:
docs/tool-comparison/chained-benchmark/Dockerfile. ricopili
~~~

Dockerfiles had to be reimplemented since the original files provided in the RICPOILI package were broken and referenced non-existing files.

Comparing with the script used for executing nf-rarevar-assoc highlights the amount and complexity of “glue” code that was necessary to write, which is not needed when using nf-rare-var-assoc. The script makes multiple bcftools and plink calls, awk and sed data format conversion calls, deals with chromosome X pseudoautosomal regions and sex, calls various RICOPILI tools and transforms its outputs. The resulting bash script is a complex pipeline and was error-prone to write.

#### C.5. Running RICOPILI + STAARpipeline

Similar to RICOPILI + nf-gwas we executed a RICOPILI + STAARpipeline comparison. This time, nothing was “loaned” from nf-rare-var-assoc so this was a fully independent association run, but it required us to create multiple scripts. We wrote:

##### Dockerfile.favorannotator

zhouhufeng/FAVORannotator publishes no Dockerfile. (60 lines of code)

##### Dockerfile.ricopili

reusing the same one from RICOPILI + nf-gwas run. (235 lines of code)

##### convert vcf to gds.R

STAARpipeline input is a GDS file, not VCF, so we had to do the conversion. (56 lines of code)

**TABLE S3:** Parameters used for dataset generation. “Domain” is the distribution the parameter was drawn from - when it is given between the curly brackets it means the values were uniformly selected from given multi-set. Repetitions in the multi-sets are on purpose, to increase the likelihood of a given value being selected.

| Parameter | Type | Domain / distribution | Description |
| --- | --- | --- | --- |
| cohort_count | int | 2000 (fixed) | Total requested cohort size. |
| cohort_proportion | float | 2.0 (fixed) | Requested control-to-case ratio, so `--simu-cc 667 1333` was passed to GCTA for every dataset. |
| simu_k | float | [0.003, 0.3], log-uniform | Disease prevalence, determines how many cases are realised. |
| simu_hsq | float | [0.001, 0.7], uniform | Heritability of the simulated trait. |
| min_af | categ. | {0, 0, 0, 0, 0, 0.001, 0.01, 0.05} | Lower bound of the allele-frequency window (`--min-af`). |
| af_range | categ. | {0.05, 0.05, 0.1, 0.2} | Width of the allele-frequency window - the upper bound passed to `--max-af` is `min_af + af_range`. |
| qual_th | int | [3, 15], uniform | Lower bound on site QUAL. |
| gq_th_filter1 | int | [3, 15], uniform | Lower bound on average genotype quality `AVG (FORMAT/GQ)`. |
| dp_min_filter1 | int | [3, 15], uniform | Lower bound on average read depth `AVG (FORMAT/DP)`. |
| dp_max_filter1 | int | [150, 300], uniform | Upper bound on average read depth. |
| min_variant_count | int | [2, 6], uniform | Minimum number of variants, after filtering, a gene must carry to be eligible as a causal gene. |
| k_genes | int | [1, 12], uniform | Number of genes selected as causal. |
| variant_fraction | float | [0.05, 0.7], uniform | Fraction of a causal gene’s surviving variants designated causal and passed to `--simu-causal-loci`. |
| max_allele_length | int | [20, 70], uniform | Maximum allele length, bounding indel size. |
| annotations_list | categ. | 2 choices: a strict expression with probability 4/5, and a no-filter with probability 1/5 | Filter on the VEP CSQ field. The strict expression matches 25 loss-of-function, missense and inframe consequence terms; the no-filter alternative is `CSQ ~ ". * "`. |

##### favorannotator csv essential.R

annotating a GDS file using the FAVORannotator. (178 lines of code)

##### genesis pcair pcrelate.R

runs GENESIS PC-AiR + PC-Relate to produce the principal components and PC-Relate GRM needed by STAARpipeline - this is actually not RICOPILI or STAARpipeline, so technically this pipeline should be named RICOPILI + GENESIS + STAARpipeline. STAARpipeline tutorials also use GENESIS PC-AiR + PC-Relate so we followed to do the same. (200 lines of code)

##### staar gene centric coding.R

running STAARpipeline gene-centric association analysis over a FAVORannotator aGDS. (357 lines of code)

##### run chain ricopili staar.sh

the main script orchestrating the whole pipeline run, executing RICOPILI, GENESIS, STAARpipeline steps, along with various bcftools and plink calls and sed/awk data conversion steps. (392 lines of code)

In addition, we provided the implementation of the results scoring, but ignoring that, to run a pipeline comparable to nf-rare-var-assoc composed of RICOPILI and STAARpipeline we had to write a total of 1478 lines of Dockerfile, R and shell code and this is exactly what a potential user of those tools faces and what is avoidable when using nf-rare-var-assoc.

We checked the performance of the composite RICOPILI + GENESIS + STAAR pipeline with and without the Saddlepoint Approximation (SPA) correction, which is meant to improve the results on imbalanced cases/-controls datasets (which our testing ones certainly were). STAARpipeline supports SPA but only for burden tests so we tested two configurations: SPA + STAAR-B (burden tests) versus no-SPA + STAAR-O (combination of burden and variance component tests). We present the aggregated metrics of those cases in Table S5.

In the end we used no-SPA + STAAR-O for our final comparison against nf-rare-var-assoc and RICOPILI + nf-gwas.

The results we obtained point to nf-rare-var-assoc being a little better than the RICOPILI + STAARpipeline combination, although the difference was not statistically significant. Also, although we didn’t systematically measure execution time, running STAARpipeline took us noticeably longer, on the order of 5-6 hours per dataset and cumulatively several days for all the datasets, even when running 5 executions in parallel (which was the maximum we could afford taking into account our memory budget of 50 GB RAM), than both nf-rare-var-assoc and nf-gwas, which both executed in roughly 10 hours for all 30 datasets.

**TABLE S4:** Parameters of nf-rare-var-assoc.

| Parameter | Figure 1A process | Description |
| --- | --- | --- |
| filter_vcf_qual_min | BCFTOOLS_VIEW_AND_FILTER2 | Variant minimum QUAL threshold. |
| filter_vcf_avg_gq_min | BCFTOOLS_VIEW_AND_FILTER2 | Minimum average genotype quality. |
| filter_vcf_avg_dp_min | BCFTOOLS_VIEW_AND_FILTER2 | Minimum average read depth. |
| filter_vcf_avg_dp_max | BCFTOOLS_VIEW_AND_FILTER2 | Maximum average read depth. |
| filter_vcf_sample_gq_min | BCFTOOLS_VIEW_AND_FILTER2 | Per-sample GQ threshold. |
| filter_vcf_sample_dp_min | BCFTOOLS_VIEW_AND_FILTER2 | Per-sample minimum DP threshold. |
| filter_vcf_sample_dp_max | BCFTOOLS_VIEW_AND_FILTER2 | Per-sample maximum DP threshold. |
| plink2_makepgen_3_options | PLINK2_MAKEPGEN_3 | Plink filtering common to REGENIE step 1 and PCA data paths. |
| plink2_makepgen_4_options | PLINK2_MAKEPGEN_4 | Plink filtering for REGENIE step 2. |
| plink2_makepgen_5_options | PLINK2_MAKEPGEN_5 | Plink filtering for REGENIE step 2. |
| plink2_write_snplist_qc_options | PLINK2_WRITE_SNPLIST_1 | Plink filtering for REGENIE step 1. |
| plink2_write_snplist_step2_options | PLINK2_WRITE_SNPLIST_2 | Plink filtering for REGENIE step 2. |
| plink2_missing_per_pheno_options | FILTER_MISSING_PER_PHENO | Plink filtering common for REGENIE step 1, step 2 and PCA data paths. |
| plink2_indep_pairwise_options | PLINK2_INDEP_PAIRWISE,<br>F_COEFFICIENT_FILTERING | Plink filtering for F-coefficient filtering and PCA data paths. |
| plink2_indep_pairwise_window | F_COEFFICIENT_FILTERING | Plink LD window settings for F-coefficient filtering data path. |
| plink2_indep_pairwise_window_pca | PLINK2_INDEP_PAIRWISE | Plink LD window settings for PCA data path. |
| plink2_king_cutoff_threshold_pca | PLINK2_KING_CUTOFF | Plink KING cutoff threshold for PCA data path. |
| regenie_step1_options | REGENIE_STEP1 | REGENIE step 1 options. |
| regenie_step2_options | REGENIE_STEP2 | REGENIE step 2 options. |

**TABLE S5:** Ablation for the STAARpipeline: SPA + STAAR-B versus no-SPA + STAAR-O. Large standard deviations are a result of vastly different difficulty of the datasets. We also executed a pairwise comparison but it also produced a non-significant difference (p-value for average precision difference came-out out as 0.67).

| Scenario | Mean average precision |
| --- | --- |
| SPA + STAAR-B | 0.1087 ± 0.140 |
| no-SPA + STAAR-O | 0.1176 ± 0.138 |

## Appendix D. Functionalities comparison

Table S6 contains the rankings of 9 tools on 23 evaluation dimensions. Grading is in the [1, 5] range, where grade 1 means the feature support is the most complete and grade 5 means it’s absent. The description of all the evaluation dimensions, grading criteria and justifications for non-obvious tool grades, can be found below.

### D.1. Inputs & formats

Which input file types the tool accepts, and whether it works with hg38 reference genome.

#### Grading criteria

1. supports VCF and popular plink formats and hg38
2. supports only plink formats or VCF, not both, but supports hg38
3. works only with other input formats, not VCF or plink, but supports hg38
4. supports VCF or plink formats but doesn’t support hg38
5. works only with formats other than VCF or plink and doesn’t support hg38

#### Individual grade justifications

##### nf-rare-var-assoc - (2)

ingests only multi-sample VCF, hg38 is the default and

works end to end (overridable)

##### montilab/nf-gwas-pipeline - (4)

ingests VCF, no PLINK option, hg38 does **not** work for gene-based tests (hg19 hardcoded in `04_gene_based.R` and example configs state hg38 is not available)

##### STAARpipeline - (3)

neither VCF nor PLINK, only an annotated GDS supported as input, hg38 works (but is the only reference genome version supported, and also hardcoded)

##### SEQSpark - (2)

ingests VCF, no PLINK option, hg38 works (“hg19” hardcoded in /`Annotation.scala` but this is not used)

##### RICOPILI - (1)

ingests both VCF and PLINK1 bed/bim/fam, hg19 is the default but hg38 does work through liftover

##### plinkQC - (1)

ingests PLINK1 bed/bim/fam and VCF (via**i** `convert_from_vcf()` from `ancestry.R`), hg38 works and is the default

**TABLE S6:** Functionality comparison of nf-rare-var-assoc to similar association pipelines and QC-focused pipelines. Dimensions manually graded 1–5 by completeness, lower = more complete. Compared tools: **a**: nf-rare-var-assoc, **b**: genepi/nf-gwas, **c**: HTGenomeAnalysisUnit/nf-pipeline-regenie, **d**: montilab/nf-gwas-pipeline, **e**: UW-GAC GENESIS analysis pipeline, **f**: STAARpipeline, **g**: SEQSpark, **h**: RICOPILI, **i**: plinkQC.

| Dimension | a | b | c | d | e | f | g | h | i |
| --- | --- | --- | --- | --- | --- | --- | --- | --- | --- |
| *Data engineering / preparation* |  |  |  |  |  |  |  |  |  |
| 1. Inputs & formats | 2 | 1 | 1 | 4 | 1 | 3 | 2 | 1 | 1 |
| 2. Normalization & prep | 1 | 5 | 5 | 5 | 5 | 5 | 4 | 3 | 4 |
| 3. Dosage & GL handling | 1 | 3 | 2 | 3 | 5 | 5 | 3 | 3 | 5 |
| 4. Genotype imputation | 5 | 5 | 5 | 5 | 5 | 5 | 5 | 1 | 5 |
| *Quality control* |  |  |  |  |  |  |  |  |  |
| 5. Variant-site QC | 1 | 3 | 3 | 4 | 3 | 4 | 1 | 1 | 3 |
| 6. Genotype-level QC | 1 | 5 | 3 | 5 | 5 | 5 | 1 | 5 | 5 |
| 7. Sample QC: missingness | 1 | 2 | 2 | 5 | 5 | 5 | 5 | 2 | 2 |
| 8. Sample QC: Het / inbreeding | 1 | 5 | 5 | 5 | 5 | 5 | 5 | 3 | 3 |
| 9. Sample QC: sex | 2 | 5 | 5 | 5 | 5 | 5 | 3 | 1 | 1 |
| 10. Ancestry-based sample exclusion | 5 | 5 | 5 | 5 | 5 | 5 | 5 | 1 | 1 |
| *Structure & relatedness* |  |  |  |  |  |  |  |  |  |
| 11. Relatedness / kinship | 2 | 3 | 3 | 1 | 1 | 2 | 5 | 4 | 4 |
| 12. Population structure (PCA) | 2 | 5 | 5 | 1 | 1 | 5 | 3 | 2 | 3 |
| 13. Long-range-LD region exclusion | 1 | 5 | 5 | 5 | 1 | 5 | 5 | 1 | 1 |
| *Annotation & association* |  |  |  |  |  |  |  |  |  |
| 14. Functional annotation & grouping | 2 | 3 | 3 | 5 | 4 | 2 | 2 | 5 | 5 |
| 15. Annotation-as-weight (continuous) | 5 | 5 | 5 | 5 | 1 | 3 | 1 | 5 | 5 |
| 16. RVAS association engine & tests | 2 | 2 | 1 | 4 | 1 | 1 | 2 | 5 | 5 |
| 17. Window-based (non-gene) RVAS | 5 | 5 | 5 | 5 | 2 | 1 | 2 | 5 | 5 |
| 18. Imbalance correction (Firth, SPA) | 1 | 1 | 2 | 2 | 2 | 2 | 3 | 5 | 5 |
| 19. LD clumping into loci | 5 | 5 | 1 | 5 | 5 | 5 | 5 | 1 | 5 |
| *Output & engineering* |  |  |  |  |  |  |  |  |  |
| 20. End-to-end automation | 1 | 1 | 1 | 1 | 3 | 4 | 2 | 3 | 4 |
| 21. Reporting & outputs | 2 | 2 | 1 | 2 | 2 | 4 | 4 | 2 | 3 |
| 22. Containerized | 1 | 1 | 1 | 1 | 1 | 1 | 5 | 5 | 5 |
| 23. Maintained | 1 | 2 | 1 | 3 | 4 | 1 | 5 | 3 | 1 |

### D.2. Normalization & preparation

VCF preprocessing: splitting multiallelic sites into separate records, left-alignment and normalization, removing duplicate records, assigning deterministic variant identifiers.

#### Grading criteria

1. all operations: multiallelic sites split + left-alignment and normalization + duplicate removal + ID assignment
2. almost all above-mentioned operations
3. two such operations
4. a single such operation
5. no VCF normalization

#### Individual grade justifications:. nf-rare-var-assoc - (1)

executes all mentioned transformations

##### STAARpipeline - (5)

VCF is not among supported inputs

##### SEQSpark - (4)

multiallelic split, but no deduplication, IDs assignment or reference-based left-alignment (but a simple trailing/leading bases trimming is executed)

##### RICOPILI - (3)

executes `bcftools norm -m-both`, left-alignment but no deduplication and no ID assignment

##### plinkQC - (4)

a multiallelic split executed for ancestry-check sub-path, not as a general preparation stage

### D.3. Dosage & genotype-likelihood handling

Whether the tool uses per-genotype likelihoods/probabilities (PL/GL/GP) or expected dosage (DS) rather than only hard genotype calls.

#### Grading criteria

1. can obtain dosage from either DS or, when it is not available, from PL/GL/GP, **and** additionally fills in missing genotype likelihoods/probabilities, which are then used to compute DS
2. same as 1 but without the filling of missing likelihood-s/probabilities
3. can use DS *or* PL/GL/GP, but cannot fallback when the field it requires is not supplied
4. hard calls only

#### Individual grade justifications:. nf-rare-var-assoc - (1)

imports the DS field when present, otherwise derives DS from genotype likelihoods (DS-from-PL, min-GQ gated), and repairs missing hom-ref PL from GQ before deriving DS - the only tool that handles both paths and fills missing likelihoods

##### genepi/nf-gwas - (3)

imports a precomputed `DS` field; hard-call fallback when DS is absent

##### HTGenomeAnalysisUnit/nf-pipeline-regenie - (2)

imports DS/HDS/GP with a genotype-probability filtering

##### montilab/nf-gwas-pipeline - (3)

consumes precomputed DS

##### SEQSpark - (3)

IMPUTE2 dosage / best-guess genotypes

##### RICOPILI - (3)

consumes imputation dosages from its own imputation step

#### D.4. Genotype imputation

Whether the tool reconstructs missing genotypes - either against an external reference panel (for example Eagle/SHAPEIT phasing followed by IMPUTE imputation) or with a custom internal model-based imputer.

##### Grading criteria

1 present
5 absent (includes tools that only consume imputed data or tools that fill missing calls with mean/major-allele)

##### Individual grade justifications:. STAARpipeline - (5)

basic filling of missing genotypes with mean/minor allele - not genotype imputation.

##### SEQSpark - (5)

basic filling of missing calls from allele frequency; can read IMPUTE2 output, but runs no imputation itself.

##### RICOPILI - (1)

reference-panel imputation: Eagle/SHAPEIT phasing then Minimac3/4 or IMPUTE4.

### D.5. Variant-site QC

Per-site filtering on call quality (QUAL), average genotype quality (GQ) and average read depth (DP), missingness rate, Hardy-Weinberg equilibrium (HWE) deviations, and minor-allele frequency/count (MAF/MAC).

#### Grading criteria

1. several filter classes with a refinement (separate threshold sets per downstream processing path, case/control-aware, or batch-aware)
2. several filter classes with a single threshold set
3. a single pass or a partial set
4. only one filter class is available at all
5. none

#### Individual grade justifications:. nf-rare-var-assoc - (1)

`QUAL` plus average per-site GQ and DP bcftools filtering. Several plink `--geno/--hwe/--mac/--maf` filterings with distinct threshold sets per downstream path (REGENIE step1 vs step2 vs common-variant for PCA path).

##### genepi/nf-gwas - (3)

A single plink2 call (`geno/hwe/mac/maf`) with one threshold set. Combining `--mind` and `--geno` plink variant&samples missingness filtering in a single call means the samples are filtered first, then variants, per plink operations order (which is not well suited for small sample size data). No QUAL, no average GQ/DP filtering.

##### HTGenomeAnalysisUnit/nf-pipeline-regenie - (3)

single filtering pass, one threshold set. No QUAL or average GQ/DP filter.

##### montilab/nf-gwas-pipeline - (4)

Missingness filtering and removing of monomorphic sites (which in virtually all valid real VCF files are already not included so this filter almost always does nothing). No QUAL, no GQ/DP, no HWE. Almost no QC at all.

##### UW-GAC GENESIS analysis pipeline - (3)

Expects that most filtering was done before (checks `FILTER == “PASS”`). Itself includes MAF and MAC filters, missingness filtering for LD pruning and GRM computation paths. No QUAL score, no average GQ/DP, no HWE filtering, no dedicated missingness filtering on the association set.

##### STAARpipeline - (4)

Trusts the `FILTER == “PASS”` flag plus a single MAF-based filter. No other variant site QC.

##### SEQSpark - (1)

Configurable expression-driven per-site filters (missingness, batch-missingness, HWE, MAF), also with possible different thresholds per batch.

##### RICOPILI - (1)

Several plink filtering stages with separate case/control HWE filtering thresholds, differential missingness (`--midi`) alongside geno/MAF filtering. No average-GQ/DP filter, but the case/control aware HWE is an advantage other tools are missing.

##### plinkQC - (3)

Three independent PLINK passes (missingness, HWE, MAF/MAC). Plink data formats carry no QUAL/GQ/DP to filter on.

#### D.6. Genotype-level QC

Setting individual genotype calls to missing when their per-genotype quality (GQ) or read depth (DP) is too low.

##### Grading criteria

1 both GQ and DP enforced
3 partial
5 no genotype QC

##### Individual grade justifications:. HTGenomeAnalysisUnit/nf-pipeline-regenie - (3)

GQ-only filtering

##### SEQSpark - (1)

expression-driven GQ + DP filtering

### D.7. Sample QC: missingness

Filtering samples with too many missing genotypes.

#### Grading criteria

1 with a refinement (recomputed per phenotype, or staged relative to the variant filters)
2 a single global missingness threshold (plink `--mind`) - this is a standard approach so we decided to grade it as 2, not 3
5 none

#### Individual grade justifications:. nf-rare-var-assoc - (1)

standard missingness filtering with additional per-phenotype filtering.

### D.8. Sample QC: Heterozygosity / inbreeding

Flagging/removing samples whose genome-wide heterozygosity (the F inbreeding coefficient) is an outlier, which signals contamination or poor quality.

#### Grading criteria

1 computed on LD-pruned genotypes with a data-driven (for example standard-deviation-based) cut-off
3 a fixed cut-off, or computed without LD pruning
5 none

#### Individual grade justifications:. nf-rare-var-assoc - (1)

standard deviation based F-coefficient outlier removal

on LD-pruned data.

##### RICOPILI - (3)

data is LD-pruned but the F-coefficient filtering is done with a user-specified threshold.

##### plinkQC - (3)

standard deviation based F-coefficient outlier removal, but without LD pruning of the input.

### D.9. Sample QC: sex

Quality control based on the sex information.

#### Grading criteria

1. reported-vs-genotype sex discordance computed and then used in some way, for example filtering, not only reported
2. sex inferred from the genotypes and then used in QC or as a covariate
3. a sex summary produced but not acted on
4. none

##### Individual grade justifications:. nf-rare-var-assoc - (2)

split-PAR *→*impute-sex, and imputed sex is used as covariates.

##### UW-GAC GENESIS analysis pipeline - (5)

PAR handling exists only to segment X for analysis, not for a sex check.

##### SEQSpark - (3)

a sexCheck summary reported (hg19 PAR) but apart from reporting not used in any way.

##### RICOPILI - (1)

check-sex discordance detection.

##### plinkQC - (1)

check-sex discordance detection.

### D.10. Ancestry-based sample exclusion

Whether the tool drops samples falling outside a target ancestry group.

#### Grading criteria

1 present
5 absent

#### Individual grade justifications:. nf-rare-var-assoc - (5)

keeps all samples; ancestry enters only as PC covari-

ates, with no ancestry-based removal - deliberate choice to retain as many samples as possible.

##### RICOPILI - (1)

smartpca iterative outlier removal drops ancestry/PC outliers.

##### plinkQC - (1)

projects samples onto 1000G reference PCs and excludes those a random-forest classifier assigns outside the target ancestry.

### D.11. Relatedness / kinship

How the tool accounts for relatedness among samples.

#### Grading criteria

1. An explicit kinship/relatedness matrix (GRM) is built by the tool itself and used as a random effect inside a mixed model for the association test.
2. Relatedness is modeled, but only partially: either (a) an explicit GRM enters a mixed-model null but must be supplied externally, not built by the tool, or (b) an in-pipeline kinship estimate (for example KING(Manichaikul et al., 2010)) is used only to select an unrelated subset for a later step, while the association test itself carries no explicit kinship term, but does handle it implicitly.
3. Relatedness is handled only implicitly, by a whole-genome/ridge regression with no explicit kinship term.
4. Related samples are removed by a hard filter - whether or not the tool itself performs an association test.
5. Not handled at all, or the kinship code is present but non-functional.

##### Individual grade justifications:. nf-rare-var-assoc - (2)

KING estimate selects the unrelated subset to be used

for PCA calculation; relatedness in the association modeled implicitly by REGENIE whole-genome regression, without filtering samples.

##### genepi/nf-gwas - (3)

relatedness modeled only implicitly inside REGENIE.

##### HTGenomeAnalysisUnit/nf-pipeline-regenie - (3)

same as nf-gwas.

##### montilab/nf-gwas-pipeline - (1)

KING *→*PC-Relate builds a real GRM fed to the GENESIS mixed-model null.

##### UW-GAC GENESIS analysis pipeline - (1)

KING-IBDseg + KING-robust*→* PC-Relate GRM (ancestry-adjusted), with a per-study median-kinship threshold refinement; the GRM enters the GENESIS mixed-model null.

##### STAARpipeline - (2)

A sparse GRM enters the GMMAT/GENESIS null model as a random effect, but STAARpipeline never builds the GRM itself - it is a required external input.

##### SEQSpark - (5)

some kinship code present but commented-out.

##### RICOPILI - (4)

Identity-by-descent (IBD) PI HAT used to hard-filter related samples from its own single-variant association (removes, does not model). From what we learned during our evaluation on simulated datasets, IBD over-calls on marker-starved, filtered exome data - at default thresholds it flagged ~67% of samples as related, against ~20% for KING on the same data, which was close to the ground truth.

##### plinkQC - (4)

IBD PI HAT hard filter (relatedness-aware max-retention pruning).

### D.12. Population structure (PCA)

How the principal components (PCs) used to correct for ancestry are produced.

#### Grading criteria

1 PCs computed in-pipeline by a method that jointly optimizes for non-relatedness *and* ancestral diversity when choosing the reference set (for example PC-AiR), with the remaining samples then projected onto the resulting axes.
2 PPCs computed in-pipeline via a simpler design: an unrelated subset is chosen by a kinship/IBD cutoff alone (no ancestral diversity optimization), PCA is run on that subset, and the remaining samples are projected onto the axes.
3 PPCs computed in-pipeline with no relatedness accommodation at all (plain in-sample PCA over all samples regardless of relatedness, or a method robust only to LD/outliers, for example an iterative outlier-detecting SVD, not to relatedness), or PCs obtained solely by projecting onto an external reference panel.
5 PRequires PCs to be supplied from outside (no in-pipeline computation at all).

##### Individual grade justifications:. nf-rare-var-assoc - (2)

LD-prune *→*KING cutoff (kinship only, no divergence term) *→*PCA on the unrelated subset *→*projection of the remaining samples.

##### genepi/nf-gwas - (5)

Requires precomputed external PCs; no in-pipeline PCA at all.

##### HTGenomeAnalysisUnit/nf-pipeline-regenie - (5)

Same as nf-gwas

##### montilab/nf-gwas-pipeline - (1)

kinship *and* ancestral-divergence-optimized partitioning.

##### UW-GAC GENESIS analysis pipeline - (1)

the same PC-AiR divergence+kinship partitioning as montilab.

##### STAARpipeline - (5)

Only consumes supplied PCs, no PCA of its own.

##### SEQSpark - (3)

Plain in-sample PCA over *all* samples - no relatedness handling present.

##### RICOPILI - (2)

EIGENSOFT smartpca on an IBD-filtered (kinship-cutoff-only, not divergence-optimized) unrelated subset; relatives can optionally be projected back.

##### plinkQC - (3)

No in-sample PCA at all - projects onto a precomputed external 1000G PC space for ancestry classification only.

### D.13. Long-range-LD region exclusion

Whether known high-LD regions are excluded before PCA/kinship so that a few loci do not dominate.

#### Grading criteria

1 present
5 absent

### D.14. Functional annotation & grouping

Whether the tool assigns functional consequences to variants (such as stop-gained, missense) and uses those consequences to define the masks - the variant sets over which a rare-variant test aggregates. Judged on several things at once: the breadth of the annotation-derived catalog (coding-only vs coding + non-coding groups; SNPs-only vs SNPs + indels; all chromosome supported vs not all) and the fact of running the annotation and variant grouping in-pipeline or not.

#### Grading criteria

1. broad catalog, annotation running in-pipeline along with masks-construction
2. annotations and masks constructed in-pipeline, but a limited catalog (for example coding-only, or SNPs-only, or not all chromosome supported)
3. annotation not run in-pipeline, but the tool accepts a per-variant consequence label and composes masks from those labels, so the composition of a severity tier is a configuration parameter
4. no consequence label in the interface - the tool accepts pre-built variant sets, so annotations reach the tests only through sets the user constructs beforehand, the tool cannot differentiate severity tiers of those sets
5. no annotation-based grouping at all

##### Individual grade justifications:. nf-rare-var-assoc - (2)

in-pipeline VEP annotation (all chromosomes, indels) and construction of masks and AAF bins, fully automated; groups into binary impact tiers and focuses on coding variants by design.

##### genepi/nf-gwas - (3)

no annotation engine of its own, but REGENIE’s mask definitions compose the tested sets from per-variant consequence labels - if they are supplied by the user.

##### HTGenomeAnalysisUnit/nf-pipeline-regenie - (3)

same as nf-gwas, user-supplied masks.

##### montilab/nf-gwas-pipeline - (5)

the ANNOVAR call is used only to annotate the results table so this never reaches the test. The pipeline provides no input through which functional annotation could select the variants tested.

##### UW-GAC GENESIS analysis pipeline - (4)

no annotation engine, and no consequence label in its interface: the aggregate test consumes a finished group file in which one group constitutes one tested set.

##### STAARpipeline - (2)

broad annotation-derived catalog, both coding and non-coding, but limited to SNPs only and doesn’t work for chromosome X.

##### SEQSpark - (2)

derives functional annotation inside the tool: an integrated RefSeq consequence engine, with masks expressed as filter expressions over the resulting keys. Extensible through dbNSFP/CADD joins, but the consequence vocabulary is coding-only.

### D.15. Annotation-as-weight, continuous

Whether per-variant continuous functional scores (for example CADD, annotation principal components) weight variants *inside* the test, rather than only sorting them into discrete groups.

#### Grading criteria

1 present for the variants the tool tests
5 present but restricted (for example applied to SNPs only, indels unweighted)
3 absent

#### Individual grade justifications

##### UW-GAC GENESIS analysis pipeline - (1)

accepts a per-variant weight column (Beta/user weights)

##### STAARpipeline - (3)

continuous aPC/CADD weighting, but applied to SNVs only (indels receive no weight)

##### SEQSpark - (1)

weights variants by a continuous INFO-key annotation

### D.16. RVAS association engine & tests

How completely the tool tests aggregated rare variants. Judged on the number of distinct test families available - collapsing/burden (CMC, WSS, variable-threshold) versus variance-component (SKAT) - on whether they are combined into an omnibus statistic (for example SKAT-O), and on whether the engine adds RVAS-specific analysis beyond a single per-set p-value. Imbalance correction and window-based testing, or the null model’s treatment of relatedness and structure, are scored separately and do not count here.

#### Grading criteria

1. both test families present, combined by an omnibus, plus a further RVAS-specific capability (conditional analysis, multi-trait testing, or combination across annotation weightings)
2. both families present and combined by an omnibus, with no further RVAS-specific extensions
3. limited test types support, but more than one
4. a single aggregation test
5. no aggregation tests available

#### Individual grade justifications

##### nf-rare-var-assoc - (2)

both burden-style, variance-components and their com-

bination available.

##### genepi/nf-gwas - (2)

same as nf-rare-var-assoc.

##### HTGenomeAnalysisUnit/nf-pipeline-regenie - (1)

same as nf-rare-var-assoc plus conditional analysis added as a pipeline feature (it is implemented in REGENIE, so other REGENIE pipelines might also have this functionality, but this requires adding one more input file, with known-association variants, which they lack).

##### montilab/nf-gwas-pipeline - (4)

GENESIS/GMMAT is capable of more, but the aggregate test is hardcoded to Burden^4^, even though the

`--method` argument lists “Burden”, “SKAT”, “fastSKAT”, “SMMAT”, “SKATO”.

##### UW-GAC GENESIS analysis pipeline - (1)

GENESIS Burden / SKAT / SKAT-O, plus SMMAT and fastSKAT, plus conditional analysis.

##### STAARpipeline - (1)

STAAR-O combines Burden, SKAT and ACAT-V across multiple annotation weightings by Cauchy combination, and MultiSTAAR extends it to multiple traits.

##### SEQSpark - (2)

Burden-style (CMC, BRV, WSS, variable-threshold), SKAT and SKAT-O available.

### D.17. Window-based (non-gene) RVAS

Whether the tool can test genomic windows or regions not tied to a gene model (fixed or data-adaptive windows).

#### Grading criteria

1. present, fixed and dynamic windows sizes
2. present, only fixed window size (dynamic window size is less important than the fact that the feature is present so grading this as 2, not 3)
3. absent

##### Individual grade justifications

4. bin/04 gene based.RL66-L68

##### UW-GAC GENESIS analysis pipeline - (2)

fixed sliding-window aggregate test.

##### STAARpipeline - (1)

fixed sliding window plus dynamic data-adaptive windows (SCANG).

##### SEQSpark - (2)

fixed sliding-window aggregate test.

### D.18. Imbalance correction (Firth, SPA)

Correcting the test when cases and controls counts are very unequal.

#### Grading criteria

1. both Firth and SPA available
2. one robust method (Firth or SPA)
3. only an alternative such as permutation/resampling
4. neither

##### Individual grade justifications:. nf-rare-var-assoc - (1)

REGENIE with both Firth and SPA.

##### genepi/nf-gwas - (1)

REGENIE with both Firth and SPA.

##### HTGenomeAnalysisUnit/nf-pipeline-regenie - (2)

REGENIE supports both Firth and SPA but the pipeline hardcoded Firth, not possible to use SPA.

##### montilab/nf-gwas-pipeline - (2)

SPA only.

##### UW-GAC GENESIS analysis pipeline - (2)

SPA only.

##### STAARpipeline - (2)

SPA only.

##### SEQSpark - (3)

a resampling-based alternative only.

### D.19. LD clumping into loci

Whether the tool reduces the associated variants to independent lead signals by LD clumping.

#### Grading criteria

1 present
5 absent

##### Individual grade justifications:. HTGenomeAnalysisUnit/nf-pipeline-regenie - (1)

clumps association results into independent loci.

##### RICOPILI - (1)

plink clumping of association results.

### D.20. End-to-end automation

How much of the path from raw input to results runs as one configured pipeline, rather than being assembled by the user from separate calls.

#### Grading criteria

1. a configure-and-run pipeline: raw input to results with no user-written glue code
2. a pipeline, but needing heavy external orchestration or cluster infrastructure to stand up
3. the major stages are each automated as discrete calls, but the user writes the glue between them (format conversions, wiring)
4. a toolkit or a library of high-level functions the user scripts into a workflow. For the purpose of this evaluation a tool such as plink or bcftools is considered lower level and would be assigned grade 5. Toolkit graded as 4 must provide operations which incorporate several steps of processing with tools graded at level 5
5. a single-purpose tool or a low-level library from which the user builds essentially the whole workflow

##### Individual grade justifications:. nf-rare-var-assoc - (1)

Nextflow pipeline, configure-and-run.

##### genepi/nf-gwas - (1)

Nextflow pipeline, configure-and-run.

##### HTGenomeAnalysisUnit/nf-pipeline-regenie - (1)

Nextflow pipeline, configure-and-run.

##### montilab/nf-gwas-pipeline - (1)

Nextflow pipeline (though older DSL1), configure-and-run.

##### UW-GAC GENESIS analysis pipeline - (3)

not one pipeline - the user runs several separate per-phase submission scripts and wires each phase’s output into the next phase’s config; each phase is itself an automated multi-step cluster driver (so more than a library), but there is no single end-to-end run.

##### STAARpipeline - (4)

an R library (its “pipeline” is a set of tutorial shell/array scripts); getting from a raw VCF to results (which we did for the empirical evaluation) required writing the data format conversions, glue code, annotation setup, a main driver script and also a Dockerfile for FAVORannotator, which was missing).

##### SEQSpark - (2)

one config drives QC/annotation/association, but requires a Spark cluster to run.

##### RICOPILI - (3)

automates QC and PCA as discrete calls, but raw VCF to results still needs hand-written plink/bcftools/awk steps between them, with no single orchestrator.

##### plinkQC - (4)

an R QC package; the user scripts the function calls into a workflow. Some operations automate execution of several plink calls under the hood.

### D.21. Reporting & outputs

Result tables, diagnostic plots, and run reports.

#### Grading criteria

1. results + diagnostics (Manhattan/QQ) + a report whose association section is itself extended (an interactive viewer, clumped loci, and/or built-in multiple-testing correction)
2. results + standard diagnostics + a report
3. a partial report (tables without plots, or a QC report with no association output)
4. raw tables/counts only
5. none

##### Individual grade justifications:. nf-rare-var-assoc - (2)

HTML report with Manhattan and QQ plots + data characteristics (DP/GQ/DS/missingness distributions) plots + PCA plot + the per-step data-flow tracking report.

##### genepi/nf-gwas - (2)

Interactive HTML Manhattan + QQ plots + per-phenotype HTML report.

##### HTGenomeAnalysisUnit/nf-pipeline-regenie - (1)

HTML report with Manhattan and QQ plots, regional plots for top N clumped loci, a separate rare-variant report with per-test-group Bonferroni thresholds.

##### montilab/nf-gwas-pipeline - (2)

HTML reports, Manhattan and QQ, a separate gene-based QQ, ANNOVAR annotation of top hits.

##### UW-GAC GENESIS analysis pipeline - (2)

Manhattan and QQ with MAC- and MAF-binned QQ panels, PCA and kinship plots, HTML reports for both the null model and the association step, LocusZoom regional plots.

##### STAARpipeline - (4)

raw p-value tables; plotting is in a separate package.

##### SEQSpark - (4)

text result tables + qc.txt counts only.

##### RICOPILI - (2)

Manhattan, QQ, forest and PCA plots compiled into a LaTeX PDF report.

##### plinkQC - (3)

rich QC reports but no association output.

### D.22. Containerized

Whether the tool ships a container image.

#### Grading criteria

1 present
5 absent

### D.23. Maintained

Recent maintenance activity.

#### Grading criteria

1. active, last commit 0-6 months ago
2. a little dated, last commit 6-12 months ago
3. dated or low activity, last commit 12-24 years ago
4. unmaintained, last commit 2-5 years ago
5. abandoned, last commit more than 5 years ago

##### Individual grade justifications:. nf-rare-var-assoc - (1)

actively developed.

##### genepi/nf-gwas - (2)

last update 10 months ago.

##### HTGenomeAnalysisUnit/nf-pipeline-regenie - (1)

last update 1 month ago.

##### montilab/nf-gwas-pipeline - (3)

DSL1 pipeline, one commit in the last year, 13 months ago.

##### UW-GAC GENESIS analysis pipeline - (4)

frozen, last commit 4 years ago but it seems the development has moved to the WDL repo.

##### STAARpipeline - (1)

last commit 2 months ago, FAVORannotator also current (2026-05-08).

##### SEQSpark - (5)

abandoned, last commit almost 7 years ago.

##### RICOPILI - (3)

last commit 16 months ago.

##### plinkQC - (1)

last commit 4 months ago.

## Appendix E.

### Additional information

Execution time on a single example dataset for the pipeline and individual steps can be seen on Figure S1.

**Figure S1:**
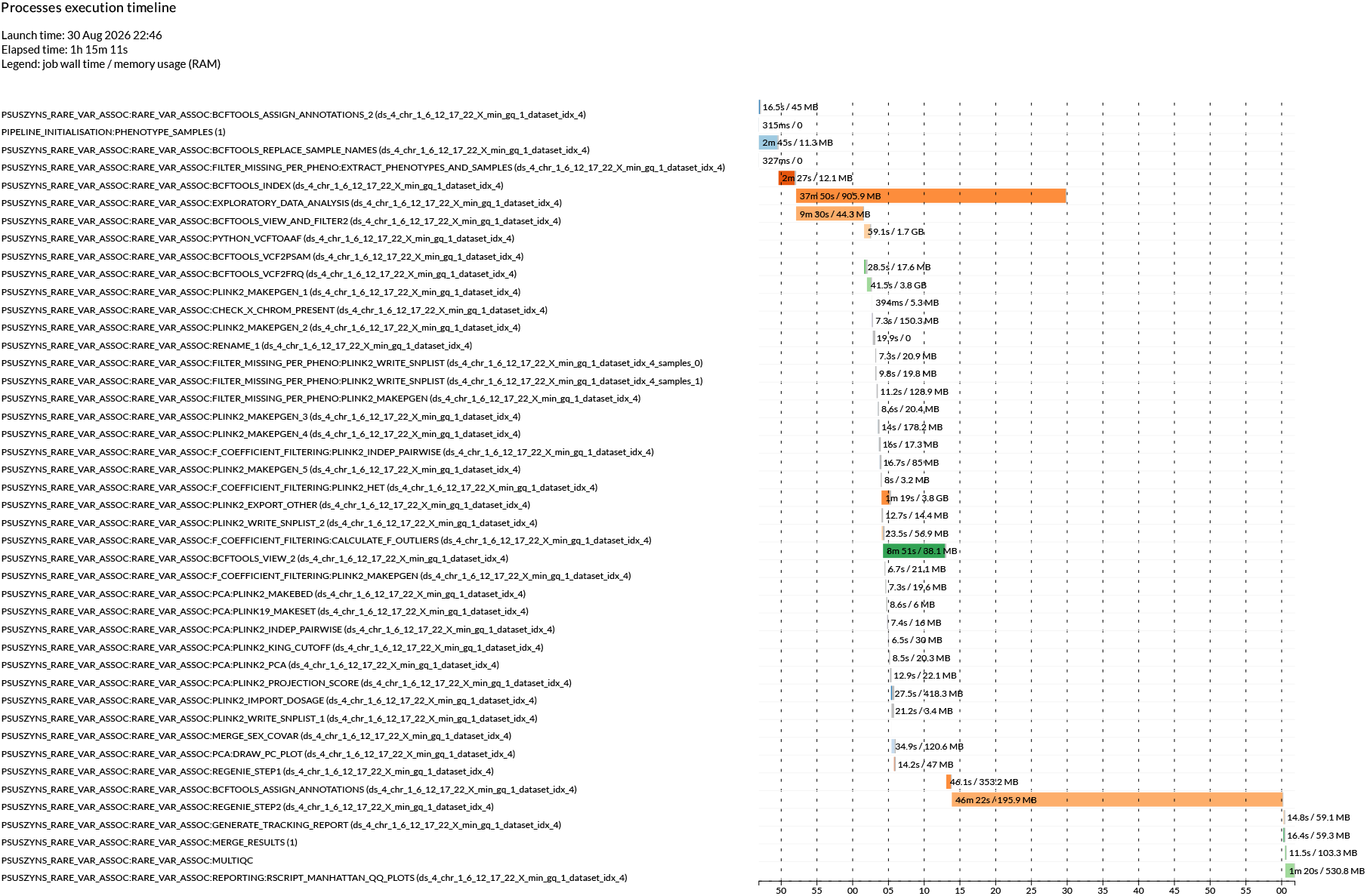
Execution timeline of the nf-rare-var-assoc pipeline for a single dataset containing exome-range limited data for chromosomes 1,6,12,17,22 and X. The most computationally expensive step is the step 2 of REGENIE, followed by the data analysis step, which however can run in parallel to other tasks so in the end it doesn’t increase the overall running time of the workflow. The other expensive steps are the bcftools filtering and indexing steps. Overall, the two REGENIE steps are responsible for 62% of the running time, highlighting the limited overhead of the whole pipeline over bare REGENIE. Note: the execution timeline omits the nested nf-prepare-vcf run (`--skip_preparation` was used) - we present the execution timeline of the nf-prepare-vcf on Figure S2. The nf-prepare-vcf pipeline must be run only once for a VCF file, after which the nf-rare-var-assoc can be run multiple times, with different settings or phenotypes.

**Figure S2:**
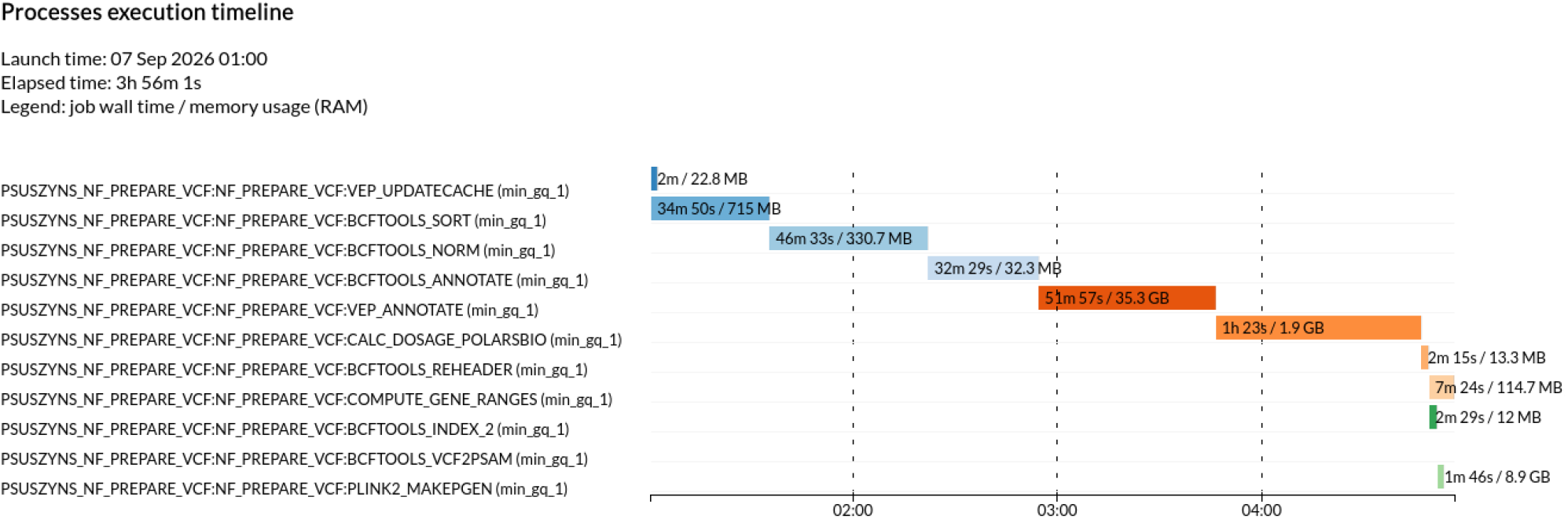
Execution timeline of the nf-prepare-vcf pipeline for a single dataset containing exome-range limited data for chromosomes 1,6,12,17,22 and X with `--calc_ds_min_gq 1` setting. VEP memory usage can be controlled with parameters such as `--buffer_size` so shown usage can be reduced. The example shows execution with a single CPU core limitation.

## Footnotes

1 https://www.cog-genomics.org/plink/2.0/formats#traw

2 Also available at https://ftp.ncbi.nlm.nih.gov/1000genomes/ftp/1000G_2504_high_coverage/working/20201028_3202_raw_GT_with_annot/

3 For more infromation read docs/tool-comparison/README.md

## Notes

### Competing Interest Statement

The authors have declared no competing interest.

